# Quantifying the individual auditory and visual brain response in 7- month-old infants watching a brief cartoon movie

**DOI:** 10.1101/610709

**Authors:** Sarah Jessen, Lorenz Fiedler, Thomas F. Münte, Jonas Obleser

**Affiliations:** Department of Neurology, University of Lübeck, Lübeck, Germany; Department of Psychology, University of Lübeck, Lübeck, Germany

**Keywords:** EEG, audiovisual, forward encoding models, temporal response function, ecologically valid stimuli, developmental neuroscience

## Abstract

Electroencephalography (EEG) continues to be the most popular method to investigate cognitive brain mechanisms in young children and infants. Most infant studies rely on the well-established and easy-to-use event-related brain potential (ERP). As a severe disadvantage, ERP computation requires a large number of repetitions of items from the same stimulus-category, compromising both ERPs’ reliability and their ecological validity in infant research. We here explore a way to investigate infant continuous EEG responses to an ongoing, engaging signal (i.e., “neural tracking”) by using multivariate temporal response functions (mTRFs), an approach increasingly popular in adult-EEG research. *N*=52 infants watched a 5-min episode of an age-appropriate cartoon while the EEG signal was recorded. We estimated and validated forward encoding models of auditory-envelope and visual-motion features. We compared individual and group-based (‘generic’) models of the infant brain response to comparison data from *N*=28 adults. The generic model yielded clearly defined response functions for both, the auditory and the motion regressor. Importantly, this response profile was present also on an individual level, albeit with lower precision of the estimate but above-chance predictive accuracy for the modelled individual brain responses. In sum, we demonstrate that mTRFs are a feasible way of analyzing continuous EEG responses in infants. We observe robust response estimates both across and within participants from only five minutes of recorded EEG signal. Our results open ways for incorporating more engaging and more ecologically valid stimulus materials when probing cognitive, perceptual, and affective processes in infants and young children.

## Introduction

Neuroimaging studies in healthy human infants are subject to severe constraints, as participants cannot follow verbal instructions, show generally short attention spans, and overall tend to be not very cooperative. As functional magnetic resonance imaging (fMRI) studies are difficult to realize in infants (Ellis & Turk-Browne, 2018), electroencephalography (EEG) continues to be the most popular method to investigate cognitive brain mechanisms in very young children and infants.

To analyze the EEG signal, most studies in infants rely on the use of event-related brain potentials (ERPs). Accordingly, most infant EEG paradigms have been optimized for the computation of ERPs: This method necessitates that a few, carefully selected stimulus conditions are repeated multiple times to elicit and average a stereotypical brain response (i.e., an ERP) that can then be compared between conditions or between individuals. This leads to experimental designs that are often (a) highly unnatural and (b) have difficulties capturing the infants’ attention for more than a few minutes.

However, in recent years and with the advent of modern computational capabilities, several new approaches to analyze EEG data have become available in adult EEG research. One such approach is the so-called “neural tracking”, which seeks to compute and assess the relationship between the recorded EEG signal and an ongoing stimulus signal. The key ideas here are, first, naturally varying, non-repetitive stimuli, often movies (Bartels, Zeki, & Logothetis, 2008; Hasson, Nir, Levy, Fuhrmann, & Malach, 2004; Nishimoto et al., 2011) or naturally spoken conversation (Broderick, Anderson, Di Liberto, Crosse, & Lalor, 2018; Ding & Simon, 2013; Fiedler, Wöstmann, Herbst, & Obleser, 2019), which have higher ecological validity and arguably engage the participant qualitatively differently than artificial, isolated stimuli (Hamilton & Huth, 2018; Huk, Bonnen, & He, 2018; Matusz, Dikker, Huth, & Perrodin, 2018). Second, a mathematical framework (usually a variant of the general linear model) that allows to either “reconstruct” features of such a natural stimulus based on the ongoing brain response (so-called backward or decoding models), or to “predict” the measured ongoing brain response from features of the stimulus (so-called forward or encoding models; Dayan & Abbott, 2001; Naselaris, Kay, Nishimoto, & Gallant, 2011).

While the use of these advanced EEG analysis approaches has become rapidly mainstream in non-human and adult human neuroscientific research, it is still rare in infant research. This is unfortunate, since they not only have yielded important new insights in adult research and are likely to offer the same potential in infant studies, but they may even provide higher gains in infancy research, which suffers from notoriously low data quality and quantity. It may for instance reduce attrition rates, as experimental designs can be optimized to be highly engaging for infant participants. Rather than presenting hundreds of repetitions of very similar stimuli, which raises the additional challenge of keeping a non-cooperative participant attending to the screen, participants can be presented with constantly changing, engaging videos in which stimuli are embedded.

Importantly, as in adult work, infant brain research has seen an increased interest in the use of naturalistic settings over the past years. Recent research has for instance demonstrated the feasibility of investigating interpersonal neural coupling in adult-infant-interactions (Leong et al., 2017) or the use of oscillatory brain responses in analyzing responses to dynamic social information (Jones, Venema, Lowy, Earl, & Webb, 2015). While dynamic, naturalistic settings and experimental paradigms yield important new insights into how brains behave and interact in real life rather than an abstract laboratory setting, they inherently pose the additional challenge of hard-to-predict and highly variant sensory input. Being able to directly relate a constantly changing input to ongoing brain responses would therefore also be crucial for the analysis of state-of-the-art ecologically valid experimental designs.

One particularly promising approach to do so is the use of multivariate temporal response functions (mTRFs), which allow us to linearly link ongoing, continuous environmental signals to simultaneously recorded brain responses in a mathematically deterministic and computationally straightforward (i.e., non-iterative) procedure. In adults, mTRFs have successfully been used to track the processing of ongoing speech (e.g., Fiedler et al., 2019) as well as ongoing and naturalistic visual input (O’Sullivan, Crosse, Di Liberto, & Lalor, 2017). Furthermore, Kalashnikova et al. (2018) used mTRFs in infants to analyze the processing of ongoing auditory speech signals, reporting a stronger cortical tracking for infant-directed compared to adult-directed speech (Kalashnikova, Peter, Di Liberto, Lalor, & Burnham, 2018).

We here demonstrate the feasibility and utility of a forward encoding modelling combined with non-repetitive complex multisensory stimulation in an infant population. We presented 7-month-old infants with a 4’48’’ long age-appropriate cartoon (one episode of the cartoon-show *Peppa Pig*) while recording the EEG. We focused our analysis on the processing of three low-level physical stimulus parameters; the auditory envelope, the motion content, and luminance. All three parameters have been amply investigated in both infants and adults and are known to elicit reliable ERP responses.

The auditory ERP response typically consists of a frontocentral P1–N1–P2–N2 sequence of responses, which can be clearly observed in adults and emerges in infancy and early childhood (see e.g., Wunderlich & Cone-Wesson, 2006, for a review). Compared to adults, infants tend to show a much less pronounced P1–N1 response, and the overall response is dominated by a broad P2 response (Wunderlich, Cone-Wesson, & Shepherd, 2006). The topographical distribution of the auditory evoked potential in infants is comparable to that of adults and characterized by a broad frontocentral distribution extending to parietal and temporal areas (Barnet, 1971).

The infant visual ERP to complex stimuli such as objects and faces comprises three main components; the Pb, the Nc, and the Slow Wave (Webb, Long, & Nelson, 2005). In particular, the Nc response, a frontocentral negativity typically observed between 400 and 800 ms after stimulus onset often linked to the allocation of attention has been amply investigated (de Haan, Johnson, & Halit, 2003; Reynolds & Guy, 2012).

If we were successful in estimating auditory and visual brain responses using a forward encoding model approach, we expect response functions comparable to classical evoked brain responses. Furthermore, since the combined use of auditory and visual regressors provides more information compared to the use of either regressor alone, we expected a more consistent and reliable response function when using auditory and visual regressors in one model.

Finally, while the predictive accuracy based on individual mTRFs computed on a subset of the available data reveals consistencies *within* subjects (Fiedler et al., 2017; Fiedler et al. 2019), a “generic” mTRF, that is, an average response function computed across participants, reveals consistencies *across* subjects (O’Sullivan et al., 2015; Mirkovic et al. 2015; Di Liberto & Lalor, 2017). Due to the limited amount of data available in the infant cohort we aimed to explore the potential benefit from relying on a generic model. Hence, we computed an averaged response function over n−1 participants and used this response function to model responses in the nth participant (i.e. leave-one-out cross validation). We directly contrasted the predictive accuracy obtained with these two approaches (individual mTRF versus generic mTRF) in the present data set.

The present manuscript thus extends previous approaches on infant mTRFs (Kalashnikova et al., 2018) by (a) using multisensory stimuli; (b) directly contrasting infant and adult responses; (c) testing a large group of infants (n=52); and (d) contrasting two different approaches to compute mTRFs, namely using a generic response function compared to an individual response function.

## Methods

### Infant participants

Fifty-two 7-month-old infants were included in the final sample (age: 213 ± 8 days [mean ± standard deviation (SD)]; range: 200-225, 24 female). Not untypical for infant studies (Stets, Stahl, & Reid, 2012), an additional 39 infants had been tested but could not be included in the final sample. Note also that directly prior to the experiment reported here, infants had already participated in a 5–10-minute-long ERP experiment on visual emotion perception (see below), further contributing to the drop-out rate since infants often became fussy or tired after the first experiment. In detail, infants were excluded because they did not watch the complete video (n=24); were too fussy to watch the video at all (n=10); did not contribute at least 100 s of artifact-free data (n=3); had potential neurological problems (n=1); or because of technical problems during the recording (n=1).

All infants were recruited via the maternity ward at the local hospital (Universitätsklinikum Schleswig-Holstein); were born full-term (38–42 weeks gestational age); had a birth weight of at least 2500 g; had no known neurological deficits; and grew up in predominantly German-speaking households. The study was conducted according to the declaration of Helsinki, approved by the ethics committee at the University of Lübeck, and parents provided written informed consent.

### Adult reference sample

In addition, we collected data from a reference sample of n = 33 adult participants. Data from n=5 were excluded due to technical difficulties during the recording (n=2) or failure to contribute at least 100 s of artifact-free data (n=3), leading to a final sample of n=28 (mean age: 50 years; range: 21–69, 16 female).

### Stimulus

As stimulus material we used one episode of the cartoon show Peppa Pig (Episode “Peppa Pig – The new car”, dubbed in German), an age-appropriate cartoon featuring a family of pigs and their daily life. Duration of the entire movie clip was 4’48’’, that is, 269 s or 6451 frames. Sound and visual parameters were not manipulated in any way.

### Procedure–Infants

After arrival in the laboratory, parents and infant were familiarized with the environment and parents were informed about the study and signed a consent form. The EEG recording was prepared while the infant was sitting on his/her parent’s lap. For recording, we used an elastic cap (BrainCap, Easycap GmbH) in which 27 Ag/AgCl-electrodes were mounted according to the international 10-20-system. An additional electrode was attached below the infant’s right eye to record the electrooculogram, which however we did not use for the present analyses. The EEG signal was recorded with a sampling rate of 250 Hz using a BrainAmp amplifier and the BrainVision Recorder software (both Brain Products).

For the EEG recording, the infant was sitting in an age appropriate car seat (Maxi Cosi Pebble) positioned on the floor. As part of a larger study, a t-shirt was positioned over the chest area of the infants. The t-shirt had either previously been worn by the infant’s mother (n=19) or by the mother of a different same-aged infant (n=14) or had not been worn before (n=19). This modulation was not of main interest to the present study and will not be analyzed or reported here in further detail.

In front of the infant (approximately 60 cm from the infant’s feet), a 24-inch monitor with a refresh rate of 60 Hz was positioned at a height of about 40 cm (bottom edge of the screen), resulting in a horizontal visual angle of approximately 27.5° and a vertical visual angle of approximately 17.6°. Left and right of the monitor, loudspeakers (Logitech X-140) were positioned and set to a comfortable level of loudness. When the infant was attending to the screen, the video was started and played without interruption until the end of the episode. The parent was seated approximately 1.5 m behind the infant and was instructed not to interact with the infant during the video. In case the infant became too fussy and started crying during the video, the video was aborted and the infant was excluded from further analysis.

Before this video presentation, infants had been presented with a series of photographs displaying happy and fearful facial expressions as part of the larger, maternal-odor study. Again, the results of this part of the study will not be further analyzed here.

### Procedure–Adults

Adult participants were presented with the same “Peppa Pig” movie after they had already participated in one of several unrelated EEG studies. They were informed about the study and signed a consent form. For recording the EEG signal, we used 64 Ag/AgCl active scalp electrodes positioned in an elastic cap according to the international 10-20-system. The EEG signal was recorded with a sampling rate of 1000 Hz using an ActiChamp amplifier and the BrainVision Recorder software (Brain Products).

Adult participants sat in a soundproof and electrically shielded chamber (Desone) in a comfortable chair approximately 1 m away from a 24-inch monitor with a refresh rate of 60 Hz on which the video was presented. Sound was presented from the same loudspeaker models used in the infant study, also positioned left and right to the screen (Logitech X-140).

### Analysis

Unless noted otherwise, the analysis protocol including the preprocessing was identical for infant and adult data. We analyzed the data using Matlab 2013b (The MathWorks, Inc., Natick, MA), the Matlab toolbox FieldTrip (Oostenveld, Fries, Maris, & Schoffelen, 2011), and the multivariate temporal response function (MTRF) toolbox (Crosse, Di Liberto, Bednar, & Lalor, 2016).

### Preprocessing

The data were referenced to the average of all electrodes (mean reference), filtered using a 100-Hz-lowpass and a 1-Hz-highpass filter, and segmented into 1-sec-epochs. To detect epochs contaminated by artifacts, the standard deviation was computed in a sliding window of 200 msec. If the standard deviation exceeds 80 mV at any electrode, the entire epoch was discarded, and if less than 100 artifact-free epochs remained, the participant was excluded from further analysis. An independent component analysis (ICA) was computed on the remaining concatenated data. Components were inspected visually by a trained coder (S.J.) and rejected if classified as artefactual (infants: 5 ± 2 components per participant [mean ± SD], range 1–10; adults: 26 ± 5, range 11–36). A 1–10 Hz bandpass filter was applied to the cleaned data. Adult data were downsampled to the infant-data sampling frequency of 250 Hz.

### Extraction of stimulus regressors

Regressors characterizing motion, luminance, and the sound envelope were extracted from the stimulus video. Exemplary excerpts of audio, luminance, and motion regressors are shown in Fig 1B.

**Figure 1.**
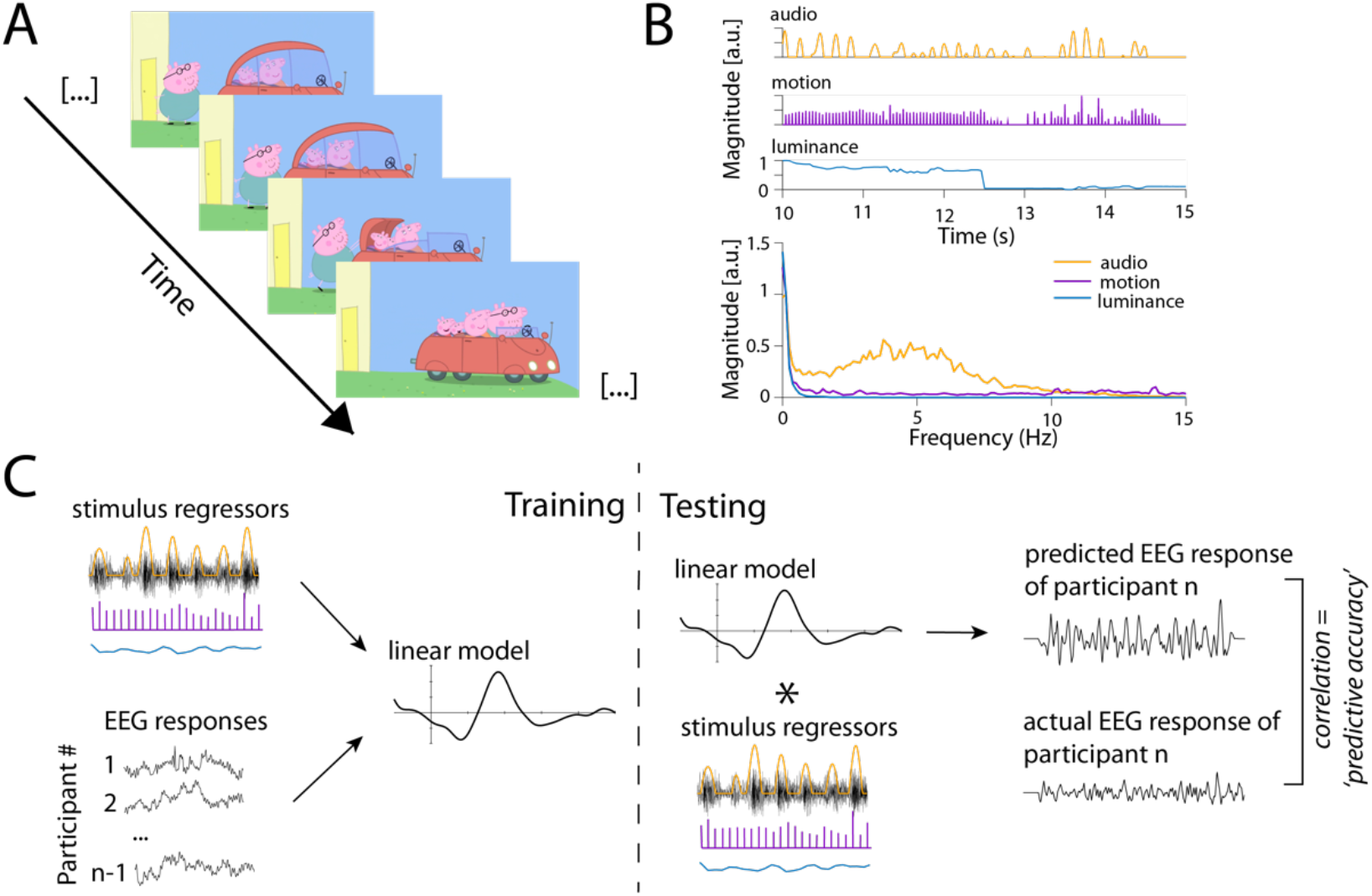
Physical properties of stimulus regressors. A) shows four exemplary stills from the movie used as stimulus material. B) shows an example of a 5-s-long stretch of the audio (orange), motion (purple), and luminance regressors (blue). Below, the frequency spectrum of the stimulus regressors is depicted; while frequencies < 10 Hz appear to be dominant in the audio regressors, no such dominance can be observed for the other regressors. C) shows an overview of the analysis approach. During training, stimulus regressors and the EEG signal of n−1 participants was used to compute a generic response function (left part). During testing, this generic response function was used to predict the EEG response of the nth participant, which was then compared to the actual EEG response of that participant. See main text for further details.

To compute a regressor of average luminance across all pixels, the weighted sum of the rgb values for each frame was computed using Matlab (Bartels et al., 2008).

To compute a regressor of average motion across all pixels, each video frame was converted to grey-scale, and the difference between two consecutive frames was computed. Then, the mean across all pixels for which this difference was larger than 10 (to account for random noise, see e.g. Jessen & Kotz, 2011; Pichon, de Gelder, & Grèzes, 2009) was computed.

To compute a regressor of sound envelope, the audio soundtrack of the video was extracted and submitted to the NSL toolbox, an established preprocessing pipeline emulating important stages of auditory peripheral and subcortical processing (Ru, 2001). The output of this toolbox resulted in a representation containing band-specific envelopes of 128 frequency bands of uniform width on the logarithmic scale with center frequencies logarithmically spaced between 0.1 and 4 kHz. To obtain the broadband temporal envelope of the audio soundtrack, these band-specific envelopes were then summed up across all frequencies to obtain one temporal envelope. Following earlier own and others’ approaches, we used the first derivative of the half-wave rectified envelope as the final audio regressor (for details see Fiedler et al., 2017). The result is a pulse-train-like series of peaks where, across frequency bands, the acoustic energy rises most steeply, reflecting “acoustic edges” such as syllable onsets.

### Stimulus parameters

As expected for a child-friendly cartoon movie, frame-to-frame fluctuations in luminance were small. On average, the change in luminance from one frame to the next was 0.35 units per frame (range 0–53, median = 0.05). Note that this deviates from previous studies where the entire dynamic range of luminance (i.e., black to white) was used to quantify the temporal response function in the adult EEG response (e.g., Lalor, Pearlmutter, Reilly, McDarby, & Foxe, 2006; Vanrullen & MacDonald, 2012) or in non-human animal electrophysiological responses (Ringach & Shapley, 2004). In contrast, the luminance-derived motion regressor yielded sizable variance, with a mean frame-to-frame change of 38 units (range 0–192, median = 36).

In sum, while variance in the luminance regressor was small, both motion and audio regressors showed considerable and promising degrees of variance.

Lastly, all regressors were downsampled (audio) and interpolated (motion, luminance), respectively, to the EEG sampling frequency of 250 Hz. In all regressors, time periods in which no EEG data was available as a result of artefact rejection during preprocessing were zero-replaced. Finally, EEG data and physical regressors were aligned and available for the linear model analysis.

### Temporal response functions (TRF)

To quantify the degree to which the EEG of 7-month-olds (as well as adults) can be expressed as a linear response to stimulus features, we used regularized regression (with ridge parameter λ) as implemented in the mTRF toolbox (Crosse et al., 2016). The key idea here is to estimate a temporal response function (TRF), that is, a set of time-lagged weights *g*, with which a regressor *s* (here, the physical stimulus features) would need to be convolved (i.e. multiplied and summed) in order to optimally predict the measured EEG response *r*.

More specifically, we used a forward encoding model approach. In a first pass, we aimed to maximize the predictive accuracy of such a model by estimating so-called “generic” models, that is, we predicted the EEG data of an *n*th participant based on a “generic” temporal response function (TRF) from *n−1* participants to the auditory or visual stimulus signal. Since changes in the EEG signal are not likely to occur simultaneously with changes in the stimulus signal but rather with an (unknown) time lag, predictions were computed over a range of time lags between 200 ms earlier than the stimulus signal and 1000 ms later than the stimulus signal.

### Choosing the optimal regularization parameter λ

To obtain the optimal regularization parameter λ for each stimulus regressor separately, as well motion and audio simultaneously, we trained the respective model on the EEG data of each participant using a variety of λ values between 10^−5^ and 10^5^. We increased the exponent in steps of 0.5, and used the resulting models to predict the EEG signal for each participant. Hence, we obtained a total of twenty different models (and predictions) based on the different λ parameters. For each of these models, we then computed the mean response function across *n−1* participants and used this response function to predict EEG response of the nth participant (i.e., *n*-fold leave-one-out crossvalidation). Finally, we computed the predictive accuracy (i.e., Pearson’s correlation coefficient *r* between the predicted EEG response and the actual EEG response) for each participant, resulting in one accuracy value for each electrode (27 for infants, 64 for adults) per participant and stimulus parameter for each λ value. For each participant, stimulus parameter, and electrode, we selected the λ value maximizing *predictive* accuracy, that is, the value for which the model yielded the highest correlation between the predicted and the actual EEG. Finally, we computed the mean regularization parameter λ value by averaging across all electrodes and participants (see Table S1). This procedure was done separately for infant and adult participants, resulting in different λ values for infants and adults.

These optimal λ parameters were used in the following to train the model, resulting in separate response functions for each stimulus parameter. For each of the three physical stimulus parameters (luminance, motion, audio) we computed a separate model. In addition, we computed a model using both, motion and audio, as regressors (“joint audio-motion model”). We chose not to include luminance in this model, as the regressor for luminance did not yield any reliable model in itself (see results).

### Evaluation of temporal response functions

For statistical evaluation of the resulting response functions, we computed a cluster-based permutation test with 1000 randomizations, testing the obtained response functions against zero. A cluster was defined along the dimensions time and electrode position, with the constraint that a cluster had to extend over at least two adjacent electrodes. A type-1-error probability of less than .05 was ensured at the cluster level (Maris & Oostenveld, 2007). Note that the number and extent of the largest clusters in the originally observed data can be compared to clusters as obtained from random permutations of the data. This constitutes the actual test at the cluster level and it protects the family-wise error at the desired type-I-error rate, here also 5 %.

In addition, to assess internal validity of our model predictions on an individual basis, we computed three different predictive accuracies per participant. First, for each participant *n*, we computed the correlation between the predicted response generated on a model trained on *n−1* participants and the actual EEG response of *n* (“generic model”).

Second, rather than relying on the generic model based on *n−1* participants, we computed an individual response function for each participant (“individual model”). To that end, 80 % of the available data for a given participant were used to train the model, and the resulting response function was then correlated with the response observed in the remaining 20 % of the data.

Third, a *permuted* or null predictive accuracy (“shifted control”) was obtained. Before calculating accuracy this way, we shifted the actual EEG response for participant *n* in steps of 2 s (in order to ensure to exceed the potential autoregressive structure of the EEG data) and computed the correlation between the shifted EEG signal and the predicted response, based on the generic model trained on *n−1* participants.

To further assess the temporal unfolding of the neural tracking, changes in predictive accuracy across time were assessed using a sliding window over a range of timelags from −200 to 1000 ms (see also Fiedler et al., 2019). In particular, we used a sliding window with a width of 48 ms (12 samples) and an overlap of 24 ms (6 samples) and computed the predicted EEG signal and its correlation with the actual recorded EEG signal for each window. Finally, we selected the time-window of maximal predictive accuracy, computed the predicted EEG signal for this optimized time-window, and directly compared the obtained results to the results obtained by the generic model based on the entire range of time lags.

Finally, we used t-tests to directly compare infant and adult response functions at each sampling point over a range of time lags between −200 ms and 1000 ms, correcting the results for multiple comparison by using the false-discovery rate procedure as suggested by Benjamini and Hochberg (1995).

## RESULTS

### Temporal response function

We computed a generic temporal response function for each stimulus regressor as well as the audio and motion regressor combined (joint audio-motion model).

We observed a clearly defined response function using the audio regressor and the motion regressor (Figure 2 and Figure S2/S3 for adults), while no clear response function could be obtained using the luminance regressor for either infants or adults (Figure S1). While Figures 2 shows the respective response functions obtained from a model which included both regressors (joint audio-motion model), comparable response functions resulted when using either of the regressors in isolation. When directly comparing the predictive accuracy obtained by the joint audio-motion model to that of the models using either regressor in isolation, we observed a higher predictive accuracy for the joint model for both, the audio and the motion regressor, in infant (joint model vs. auditory only: *t*(51) = −4.4, *p*<.001; joint model vs. motion only: *t*(51) = −7.77, *p*<.001). In the adult data, this was only the case for the contrast between the joint model and the motion-only model (joint model vs. auditory only: *t*(27) = −.13, *p* = .20; joint model vs. motion only: *t*(27) = −8.44, *p*<.001).

**Figure 2.**
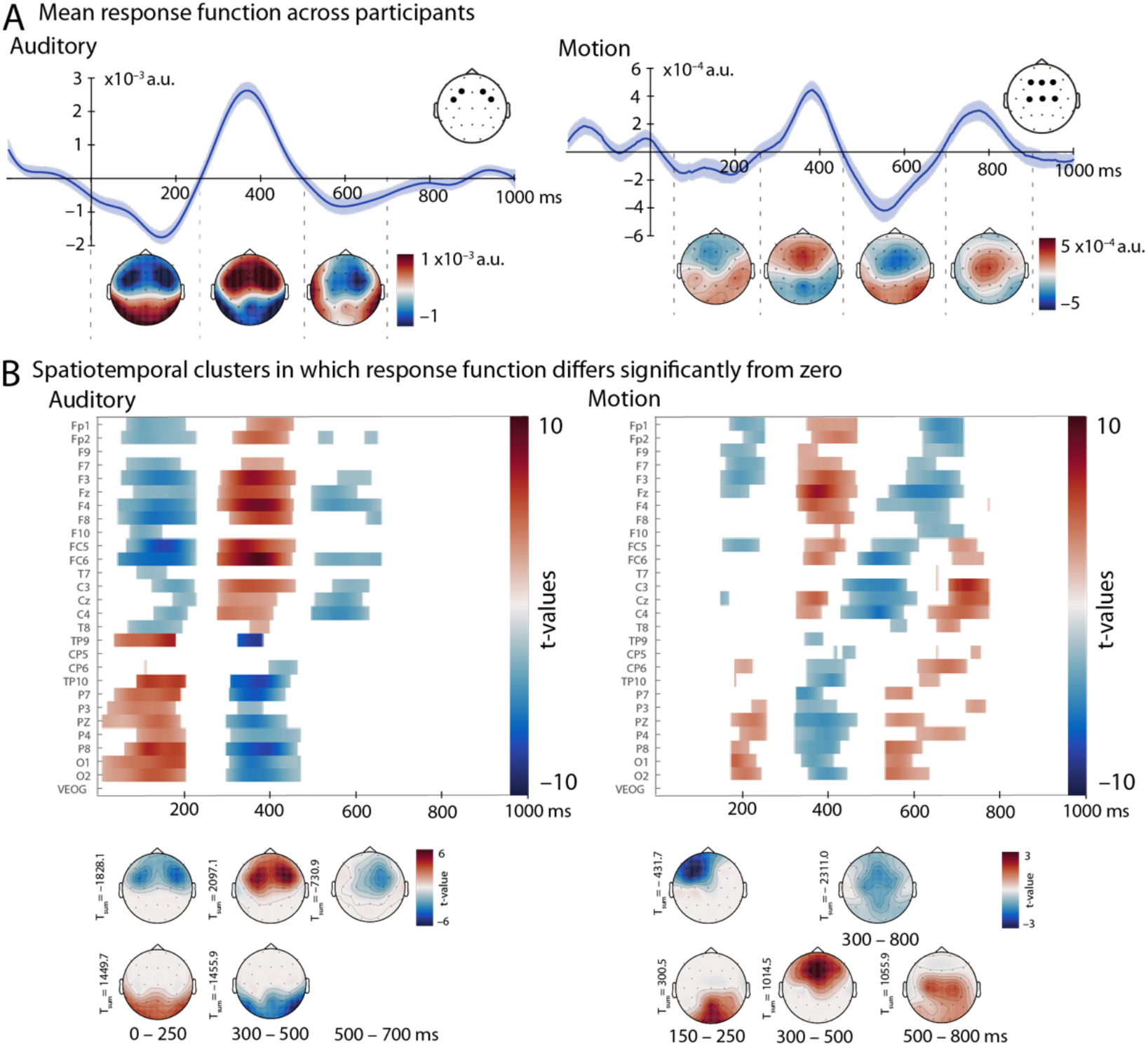
Response function (using motion and audio regressor simultaneously) for infant participants. The left panel shows the auditory response function, while the right panel shows the motion response function. A) depicts the mean mTRF (mean ± SEM) computed across all participants, averaged over FC5, FC6, F3, and F4 (auditory) respectively F3, Fz, F4, C3, Cz, and C4 (motion) with the included electrodes marked by black dots. Electrodes were chosen based on visual inspection of the topography and the cluster test (B). Topographic representations for are shown 0–250 ms, 250–500 ms, and 500–700 ms (auditory) and 50–250 ms, 250–450 ms, 450–700 ms, and 700–900 ms (motion). B) displays the results of the cluster-based permutation test, comparing the response function shown in A) to zero. Positive deviations are displayed in red, while negative deviations are shown in blue. In the bottom part of B), the same clusters as in the top part of B) are shown as topographic distributions, along with the summed t-value across the cluster.

Interestingly, while a clearly defined response was visible for both, the audio and motion regressor, the amplitude of the response function for the motion regressor was much smaller compared to the amplitude of the audio response function.

### Cluster-based permutation test

We computed a cluster-based permutation test comparing the temporal response function obtained using the motion, luminance, and audio regressor as well as the motion and audio regressor simultaneously. We did not observe any significant cluster using the luminance regressor for either infants or adults. In contrast, we did obtain multiple significant clusters, indicating a positive or negative deviation from zero, for the motion and audio regressor, both when included separately as well as in combination (see supplementary material for a full list of the results of the cluster-based permutation test using audio and motion regressor separately as well as in combination for infants and adults, Figure 2B for infant results and S2B/ S3B for adult results). The resulting clusters confirm the deflections observed in the auditory and motion response function (Figure 2A).

### Comparing the infant and adult brain responses

When comparing infant and adult response functions (Figure 3), similarities as well as striking differences emerge. Overall, amplitudes of the response functions are comparable for infants and adults, both showing the already mentioned larger amplitudes for audio regressors and smaller amplitudes for motion regressors. For both, infants and adults, the auditory response function is marked by a prominent frontocentral positivity (250–500 ms for infants, 300–450 ms for adults). While this response appears to be slightly longer for infants, overall, both latency and topography indicate a comparable response for infants and adults. In contrast, the infant auditory response function lacks a second, earlier and more central positivity, which can be observed between 150 and 250 ms in the adult auditory response function.

**Figure 3.**
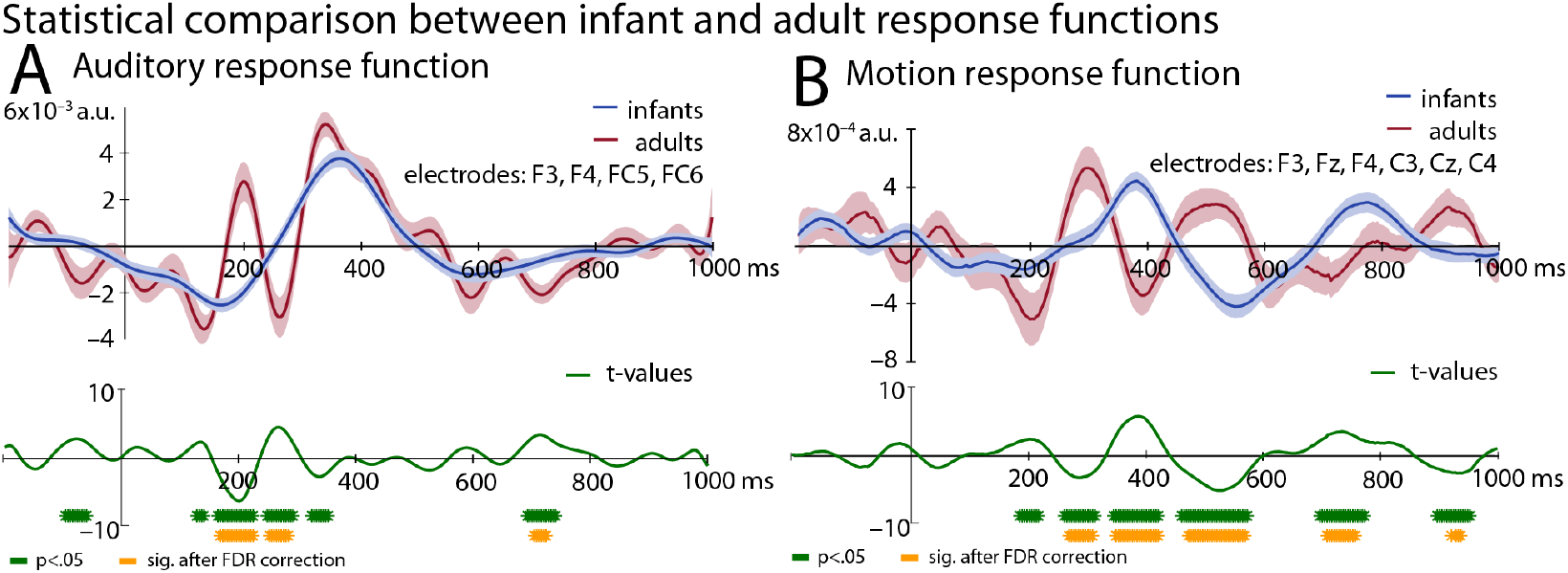
Comparison of infant and adult response functions. Mean mTRF for infant (in blue) and adult (in red) participants are shown for the audio regressor (A) and the motion regressor (B). The infant response functions and topographical representations are identical to those shown in Fig 2A and 3A for audio and motion regressors, respectively. Responses are averaged across the same electrodes for adults and infants, namely FC5, FC6, F3, and F4 for A) and F3, Fz, F4, C3, Cz, and C4 for B). The topographic representations of adult responses correspond to those in the supplementary material, namely 50–150 ms, 150–250 ms, 250–300 ms, and 300–450 ms for A) and 50–250 ms, 250–350 ms, 350–450 ms, 450–550 ms, and 550–800 ms for B). The graph in the bottom part depicts t-values of the mean infant – adult difference. Periods in which t-tests resulted in a p value <.05 are marked by green asterisks, periods in which these tests survived correction for false discovery rate are marked by orange asterisks.

For the motion response function, both infants and adults show two frontal / frontocentral positivities (250- 450 and 700–900 ms for infants and 250–350 and 450–550 ms for adults). Hence, infants and adults show a comparable response, though the infant response appears to be much slower and less temporally modulated.

When statistically comparing infant and adult responses directly, significant differences can be observed at a time lag of around 200 ms and 300 ms in the auditory response function (Figure 3A). In contrast, the motion response function differs between infants and adults at multiple lags (Figure 3B).

### Temporal unfolding of predictive accuracy

To further investigate the temporal unfolding of the response functions, we computed the predictive accuracy (i.e., the correlation between the predicted and the actual EEG response) using a sliding window of 48 ms (Figure 4). Both, infants and adults, show the highest predictive accuracy at time lags between 200 and 400 ms (“optimized time windows”). However, when using only these optimized time windows to compute the predictive accuracy, a different pattern between infants and adults emerges. In adults, predictions based on this optimized time window yielded a higher predictive accuracy compared to the generic model encompassing the entire range of time lags (*t*(27) = −6.51, *p*<.001). In infants, however, predictions based on this optimized time window yielded a lower predictive accuracy compared to the generic model encompassing the entire range of time lags (*t*(51) = 7.39, *p*<.001).

**Figure 4.**
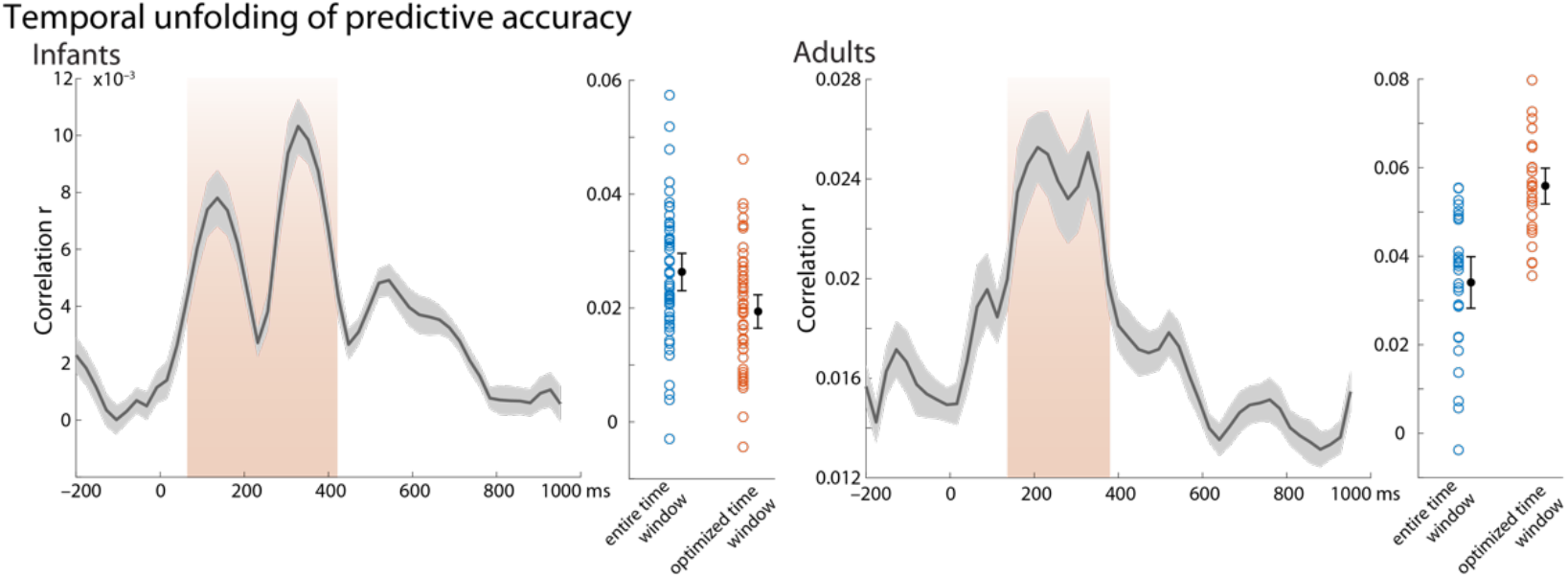
Temporal unfolding of predictive accuracy. Depicted in gray is the correlation between predicted and actual EEG in sliding time-windows of 48 ms (with 24 ms overlap, left panel: infants, right panel: adults; mean ± SEM). The area marked in orange denotes the time range used as optimized time window. Predictive accuracy was computed over the optimized time window and is shown by orange circles (next to the predictive accuracy yielded by the entire time window in blue for comparison; mean accuracies with 95 % confidence intervals are shown in black).

### Generic vs. individual response functions

The results discussed above rely on a generic model, that is, an average TRF computed based on data from *n*−1 participants was convolved with the *n*th participant’s data in order to obtain an predicted EEG trace for this subject (see Di Liberto & Lalor, 2017). An alternative approach (and in fact preferable, if enough data for per subject is available; e.g., Fiedler et al., 2019; O’Sullivan et al., 2017) computes an individual model based on a subset of an individual’s data and compare the resulting predictions to the remaining data.

As expected, individual models showed a larger variance compared to the generic model (Figure 5–7; see S4 and S5 for data from adult participants), but both, generic model and individual model result in correlations clearly above zero (with the exception of luminance, where no reliable prediction was possible for either mode, see Figure 5C; results of the t-tests against zero for the joint-model depicted in Figure 7: generic model_infant_: *t*(51) = 15.72, *p*<.001; generic model_adult_: *t*(27) = 11.49, *p*<.001; individual model_infant_: *t*(51) = 5.60, *p*<.001; individual model_adult_: *t*(27) = 8.33, *p*<.001).

**Figure 5.**
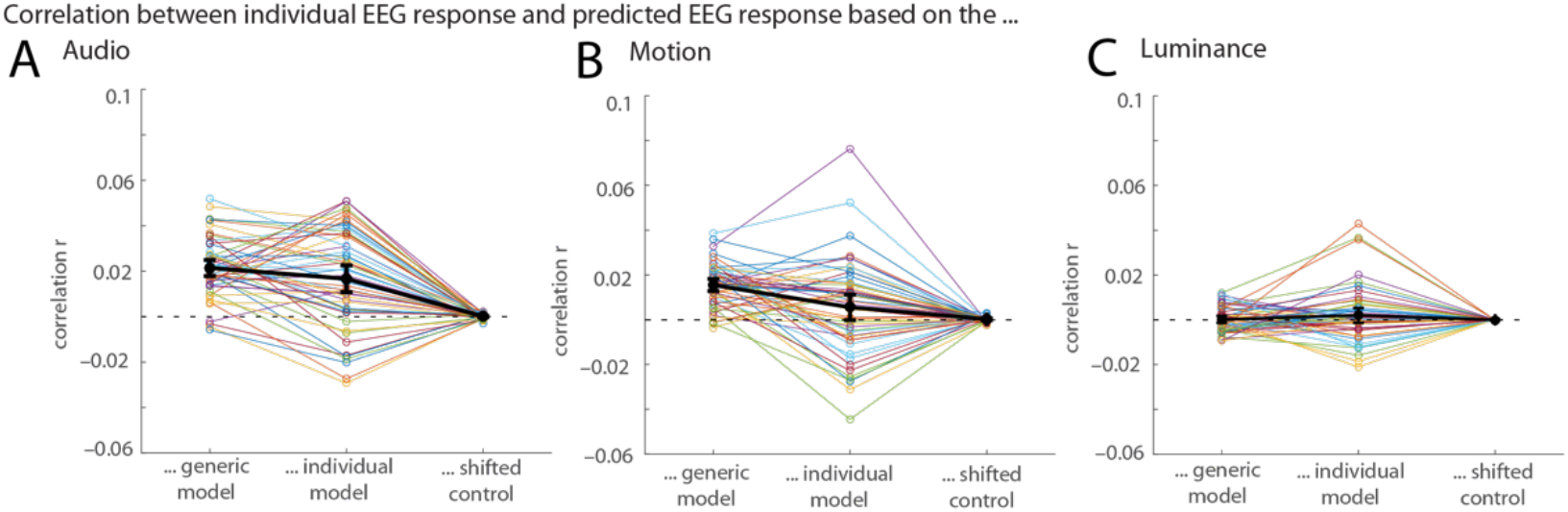
Predictive Accuracy (r) between model and EEG response for infant participants. The recorded individual EEG response was correlated with three different parameters using Pearson’s correlation coefficient for the audio regressor (A), motion regressor (B), and luminance regressor (C). On the left, the correlation between the recorded EEG responses of participant n and the response predicted by the generic model based on the remaining n−1 participants is shown for each participant. In the middle, the correlation between the model trained on the first 80 % of the data available for each participant and used to predict the remaining 20 % from that participant and the actual EEG response recoded from that participant is shown. The right column shows the correlation between the prediction generated by the generic model and the recorded EEG data shifted in a circular way in steps of 2 s as a control condition (averaged over all possible shifts). Correlations are shown for each infant participant (in colors) as well as the mean correlation with 95% CI (confidence interval) across all participants (in black).

When both, audio and motion regressor were included (Figure 6), the generic model resulted in a higher correlation compared to the individual model for infant participants (*t*(51)=3.76, *p*<.001); 37 participants showed a higher correlation with the generic model while only 15 participants showed a higher correlation with the individual model (Figure 6, right panel). Nevertheless, both approaches provide converging results, as suggested by a significant positive correlation between the predictions obtained the generic model and those obtained with the individual model (*r*=.36, *p*=.009; infants, joint regressor). When using only the motion regressor (Figure 5B), the correlations were also higher for the generic compared to the individual model (*t*(51)=3.50, *p*<.001), while for the auditory regressor (Figure 5A), this difference was less pronounced (*t*(51)=1.82, *p*=.07).

**Figure 6.**
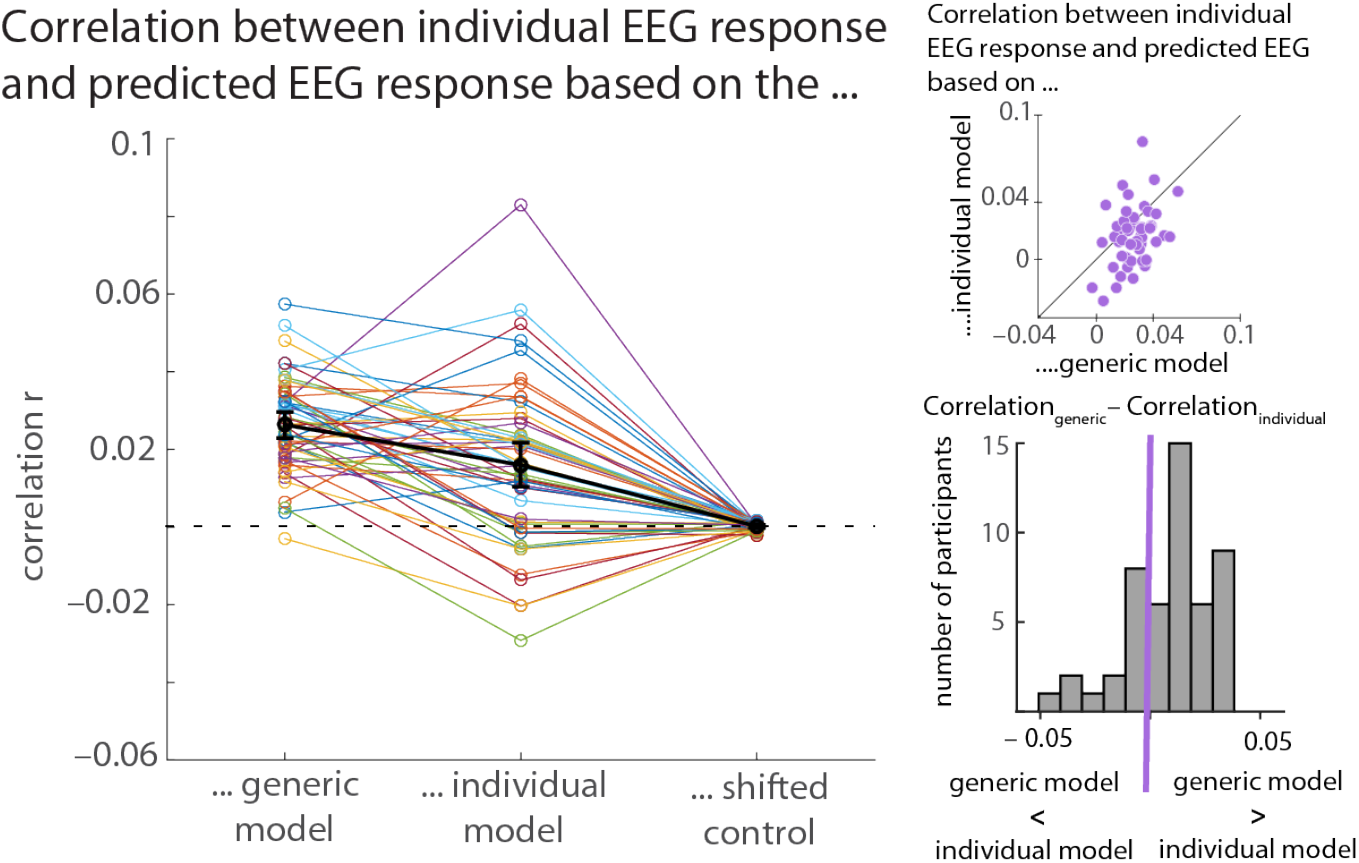
Predictive accuracy for observed EEG response for infant participants in a joint audio-motion model. The left part of the figure shows the correlation (based on Pearson’s correlation coefficient) between the recorded EEG signal and the EEG responses predicted based on the generic model (left column), the individual model (middle column), and a shifted control condition (right column, see text). The two plots on the right hand visualize a comparison between the generic and the individual model. In the top plot, each purple dot indicates the difference between the correlation with the generic model and the correlation with the individual model. Hence, a purple dot in the right bottom part of the graph indicates an individual with a higher correlation for the generic compared to the individual model, while a purple dot in the top left part indicates an individual with a higher correlation for the individual compared to the generic model. Dots on the 45° line would indicate both models to perform equally well in a given participant. The bottom plot displays the same information in a bar graph; individuals having a higher correlation for the generic model have a positive difference and hence fall to the right of the zero-threshold marked in purple while those with a higher correlation for the individual model have a negative difference and fall to the left of the zero-threshold.

**Figure 7.**
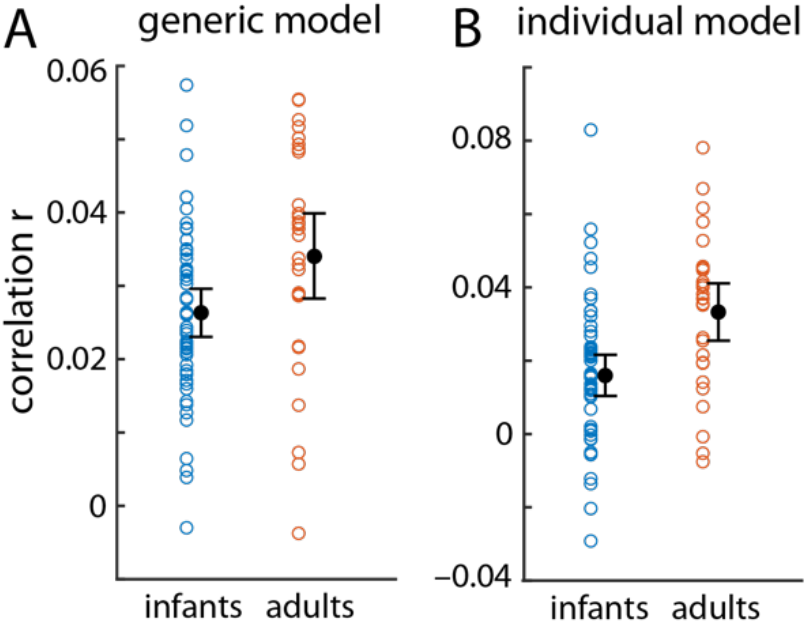
Predictive accuracy for infants and adults in a joint audio-motion model. A) shows the individual correlations using Pearson’s correlation coefficient for infants (blue) and adults (orange) using the generic model. B) shows the individual correlations using Pearson’s correlation coefficient for infants (blue) and adults (orange) using the individual model. Mean accuracies with 95 % confidence intervals are shown in black.

As a control analysis, a generic model using temporally shifted (i.e., purposefully misaligned) versions of the actual EEG signal (1,000 iterations) did yield substantially lower predictive accuracy values.

Finally, we directly compared the use of the generic compared to the individual model between infants and adults (Figure 7). For both, the generic and the individual model, the adult group showed a higher predictive accuracy (generic model: *t*(78) = −2.46, *p*=.016; individual model: *t*(78) = −3.54, *p*<.001).

## Discussion

We here investigated the use of a variant of forward encoding models (multivariate temporal response functions, mTRFs) to analyze infant brain responses to a continuous complex audiovisual stimulus, namely a 5-minute cartoon movie. We observed clearly defined response patterns to both the auditory as well as the motion content, but no predictive response function for changes in luminance was found.

Our results demonstrate that the simultaneous acquisition of individual brain responses to different sensory modalities is possible in the infant brain, opening new avenues for ecologically valid multisensory research paradigms in developmental neuroscience. Furthermore, our results suggest that a generic model derived from a larger set of unrelated infant data is as good or slightly better compared to an individual model in predicting the individual brain response, especially in cases where only limited data is available. This points to the further utility of such an approach in developmental and at-risk populations.

### Motion and Audio

For both, motion and audio information in the cartoon movie, we found a clearly defined response in both infants and adults. The observed responses are largely consistent with patterns typically reported in more traditional event-related brain potentials. In particular, the frontocentral negativity between 450 and 700 ms observed in the infant brain responses linked to the motion regressor corresponds in timing, shape, and topography to the Nc component, an infant ERP component that can routinely be observed in visual paradigms and has been linked to attention allocation (Webb et al., 2005). Furthermore, the bifocal frontal positivity observed in the infants’ brain response linked to the auditory envelope shows a strong similarity to the commonly reported P2 response in infant auditory brain responses (Wunderlich et al., 2006), though the peak can be observed around 400 ms and therefore somewhat later than a typical P2 response.

Similar comparisons can be drawn for observed response functions in the adult group. The auditory response function is characterized by an P1-N1-P2-N2 pattern, which resembles the typical adult auditory onset response (Picton, 2013). As in the infants, the response appears to be slower though, with the first positivity peaking around 200 and the second around 350 ms. The topographical distribution, namely a broad frontocentral activation, is consistent with typical auditory ERP responses (Picton, 2013). Importantly, the auditory response function also resembles the pattern that has been reported in prior studies investigating neural tracking of attended auditory information in adults (e.g. Fiedler et al., 2019).

The adult motion response function shows a pattern characterized by two positivities, one peaking around 300 ms, the other around 500 ms. The topographical distribution is again primarily frontocentral, though a positivity around 400 ms can been seen at posterior and occipital electrodes (see Figure 3 and Supplementary Figure S3). Prior comparable studies investigating the processing of continuous naturalistic stimuli have also reported a response profile characterized by two positivities, although at shorter latencies and with primarily occipital distributions (see e.g. O’Sullivan et al., 2017).

The direct comparison of infant and adult brain responses (Figure 3) may provide insight into developmental changes. In response to the auditory envelope, both infants and adults show a prominent frontal negativity peaking around 400 ms. Notably, however, the adults show an additional central positivity around 200 ms, which is missing in the infant response. This corroborates and replicates known developmental changes commonly observed in auditory evoked responses when comparing infants and adults (Wunderlich & Cone-Wesson, 2006). Considering the motion response, the correspondence between infant and adult response is less straight-forward. While the adult response is characterized by two frontocentral positivities, one peaking around 300 ms and the other around 500 ms, the infant response is dominated by one frontocentral peak around 400 ms.

Importantly, we used both, generic response functions as well as individual response functions to predict the EEG signal. When using both, the motion and the auditory regressor, performance was significantly better for the generic compared to the individual model. When using only the auditory regressor, the same pattern was visible but the difference only marginally significant. Note, however, that both, generic and individual models generated predictions that were significantly above chance level. This demonstrates two important things. First, five minutes of EEG recording are sufficient to compute reliable models, both on an individual level as well as across participants as a generic model. This is not only true for EEG data obtained from healthy adults but also for data obtained from populations providing notoriously noisy signal, such as infants. Second, brain responses across participants, both infants and adults, are sufficiently similar to generate a model that can successfully predict a new infant’s brain response, yielding even better outcomes compared to the individual model.

### Limitations and future studies

The present study provides an important step and proof of feasibility for using mTRFs to analyze infant EEG data in response to complex and dynamic audiovisual stimulus material. This offers a whole host of new possibilities in the investigation of infant’s brain responses in their natural environment.

One important advantage of using mTRFs compared to classical ERPs approaches is the possibility of using more complex and naturalistic experimental settings. These might include the processing of continuous auditory signals (as demonstrated by Kalashnikova et al., 2018), or audiovisual video material, as in the present study. In contrast to ERPs, which require the repetitive presentation of short stimuli, with mTRFs, it is possible to investigate the brain’s entrainment over a longer period of time and in a much closer approximation of natural settings. Taking this last aspect a step further, mTRFs thereby also offer a potential approach to analyze brain responses in live interactions in which the live input the infant receives (or also the infant’s responses) are recorded and used as a regressor in the subsequent analysis. Such an approach would provide an important tool in investigating the neural bases of social interactions, a topic that has received increasing interest in recent years also in developmental neuroscience (see e.g., Leong et al., 2017; Wass et al., 2018).

From a research-design point of view, the use of continuous stimulation (in contrast to short repetitive stimuli needed for ERPs) also allows for the design of more engaging experiments. This is particularly important for populations characterized by short attention spans and overall low compliance, such as young children, but also patient groups. In addition, such participant groups are often also characterized by a low signal-to-noise ratio in the EEG signal and only limited data availability; here, we demonstrate that 5 minutes of recording are sufficient to compute reliable responses, highlighting a further practical advantage of using mTRFs to analyze EEG data in special populations.

While the present study provides an important lead, it also raises several new questions. One important feature of the present study is that we used the unmanipulated cartoon video material. While this makes for an ecologically valid and easy-to-obtain stimulus, it comes with the caveat of a lack of control for stimulus properties.

Notably, while we did observe a clear-cut response to the motion and the auditory regressor, we did not find a reliable response to the changes in luminance. The most likely explanation for this discrepancy is the lack in variance in the luminance content. While the motion and the auditory regressor showed large-amplitude changes throughout the video (e.g., average motion change between frames = 38 units), average luminance of this cartoon movie remained fairly constant (average luminance change between frames = 0.35 units). Previous studies targeting neural responses to luminance change (in adults) typically used considerably more pronounced black-white contrast (Lalor et al., 2006; Vanrullen & MacDonald, 2012). Hence, the luminance changes in the stimulus material were likely too small to elicit any robust change in brain response. Future studies explicitly varying the luminance content are therefore necessary to investigate the applicability of mTRFs to other visual stimulus parameters in infants.

Also, we operationalized motion as change in pixel from one frame to the next. This means that the motion regressor not only reflected the actual motion of the objects and persons depicted in the video but also cuts in the video. For the present purpose, we did not differentiate between these two possibilities of motion. Furthermore, in the present set-up, we did not control for degree to which individual infants constantly attended visually to the screen; future studies combining EEG recordings with eye tracking might therefore further improve the predictive accuracy of the visual models.

Building upon the present results, a next step would therefore be to purposefully manipulate such parameters. By using stimulus material designed to encompass a larger variance in luminance and/or no cuts in the video, it should for instance be possible to observe brain response to changes in luminance and motion responses that can be clearly linked to actual motion rather than video cuts. Such an approach could for instance provide valuable new insights into the processing of biological motion (Marshall & Shipley, 2009; Reid, Hoehl, & Striano, 2006).

Furthermore, in the present study, we did not contrast different conditions, neither within infant nor between different groups of infants. Having demonstrated the feasibility of using encoding models to model brain responses for this type of complex audiovisual stimuli, the next step would certainly be to utilize this approach to investigate differences in processing between (a) different types of stimulation or (b) different groups of infants.

## Conclusion

The present data demonstrate that forward encoding models based on the multivariate temporal response function (mTRF) pose a valuable and versatile tool in quantifying and disentangling complex audiovisual brain responses and the according perceptual processes in infancy. Our results open way for applications to a variety of research areas not only in early development, but also in other special populations characterized by short attention spans and low cooperativeness, including research in severely impaired neurological patients. New paradigms could not only entail complex multisensory perception, but extend to dynamic social interactions. As such, mTRF approaches to infant data analysis will allow developmental researchers to devise more engaging and thereby more easily applicable experimental set-ups for infancy research.

## Acknowledgements

We thank all the families for participating, Leonie Emmerich, Aylin Ulubas, Franziska Scharata, and Anne Hermann for help with the data acquisition, and the German Research Foundation (DFG) for funding to SJ (JE 781/1-1 & 2). Three reviewers provided valuable and constructive feedback.

## Data accessibility

Data will be made available upon publication on Open Science Framework (OSF).

