## Supplementary Material for "Quantifying the individual auditory and visual brain response in 7- month-old infants watching a brief cartoon movie"

#### Tables

Table S1. Lambda parameters l obtained for infants and adults. 10^l

| | Infants | Adults |
|---|---|---|
| Audio | 0.1247 | –0.2852 |
| Motion | –0.7840 | –0.7501 |
| Luminance | 0.2201 | –0.0260 |
| Audio and Motion | –1.9190 | –2.4870 |

Table S2. Correlation r (Pearson's correlation coefficient) between number of available data points and correlation between individual MTRF trained on 80 % data and tested on 20 % data.

| | Infants | Adults |
|---|---|---|
| Audio | –.0902 | .0928 |
| Motion | –.1325 | –.1887 |
| Luminance | .0684 | –.0940 |
| Audio and Motion | –.0885 | –.0396 |

Table 3. Results of cluster-based permutation test (note that for the regressor luminance, neither infants nor adults showed any significant clusters)

| Infant | Adult |
|---|---|
| **Audio** | |
| $p<.001$, $T_{sum}= -1840.7$ | $p<.001$, $T_{sum}= -5683.7$ |
| $p<.001$, $T_{sum}= -1469.5$ | $p=.004$, $T_{sum}= -2387.8$ |
| $p<.001$, $T_{sum}= -772.5$ | $p=.0140$, $T_{sum}= -1370.5$ |
| $p<.001$, $T_{sum}= 2120.2$ | $p=.0160$, $T_{sum}= -1281.2$ |
| $p<.001$, $T_{sum}= 1525.0$ | $p<.001$, $T_{sum}= 6385.3$ |
| | $p<.001$, $T_{sum}= 3294.6$ |
| | $p=.0140$, $T_{sum}= 1254.3$ |
| **Motion** | |
| $p<.001$, $T_{sum}= -2767.1$ | $p<.001$, $T_{sum}= -1892.3$ |
| $p=.002$, $T_{sum}= -553.7$ | $p<.001$, $T_{sum}= -1820.5$ |
| $p<.001$, $T_{sum}= 1295.4$ | $p<.001$, $T_{sum}= 3417.0$ |
| $p<.001$, $T_{sum}= 1225.4$ | |
| $p=.005$, $T_{sum}= 455.5$ | |
| **Audio (using audio and motion as regressors)** | |
| $p<.001$, $T_{sum}= -1828.1$ | $p<.001$, $T_{sum}= -3912.9$ |
| $p<.001$, $T_{sum}= -1455.9$ | $p<.001$, $T_{sum}= -2367.1$ |
| $p<.001$, $T_{sum}= -730.9$ | $p=.0120$, $T_{sum}= -1488.1$ |
| $p<.001$, $T_{sum}= 2097.1$ | $p=.0130$, $T_{sum}= -1422.4$ |
| $p<.001$, $T_{sum}= 1449.7$ | $p=.0170$, $T_{sum}= -1186.7$ |
| | $p<.001$, $T_{sum}= 6094.7$ |
| | $p<.001$, $T_{sum}= 3189.5$ |
| | $p=.0100$, $T_{sum}= 1275.4$ |

| Motion (using audio and motion as regressors) | |
|---|---|
| p<.001, $T_{sum}$= -2311.0 | p<.001, $T_{sum}$= -1556.9 |
| p=.002, $T_{sum}$= -431.7 | p=.0280, $T_{sum}$= -792.4 |
| p<.001, $T_{sum}$= 1055.9 | p=.0490, $T_{sum}$= -705.8 |
| p<.001, $T_{sum}$= 1014.5 | P<.001, $T_{sum}$= 2922.2 |
| p=.005, $T_{sum}$= 300.5 | |

### Figures

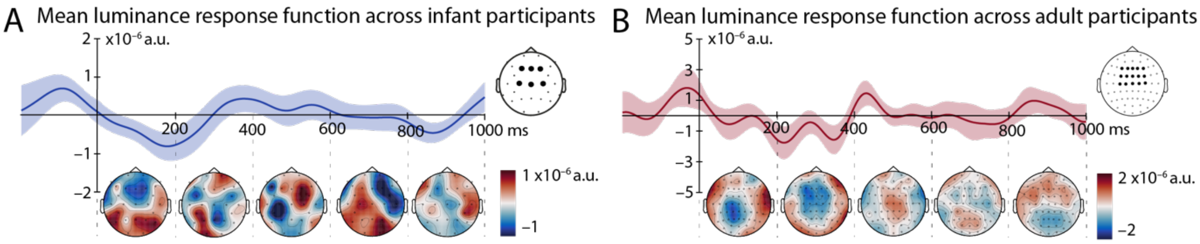


*Figure S1. Luminance response function for infant and adult participants.* A) and B) shows the generic model (mean ± within-subject SEM) computed across all participants (A: infants; B: adults), averaged over F3, F1, Fz, F2, F4, FC3, FC1, FCz, FC2, FC4, C3, C1, Cz, C2, and C4, and topographic representations for 0 – 200 ms, 200 – 400 ms, 400 – 600 ms, 600 – 800 ms, and 800 – 1000 ms with electrodes included in the above-shown average marked by black dots. Note that, unlike for auditory and motion response functions, no significant clusters were found in the cluster-based permutation test.

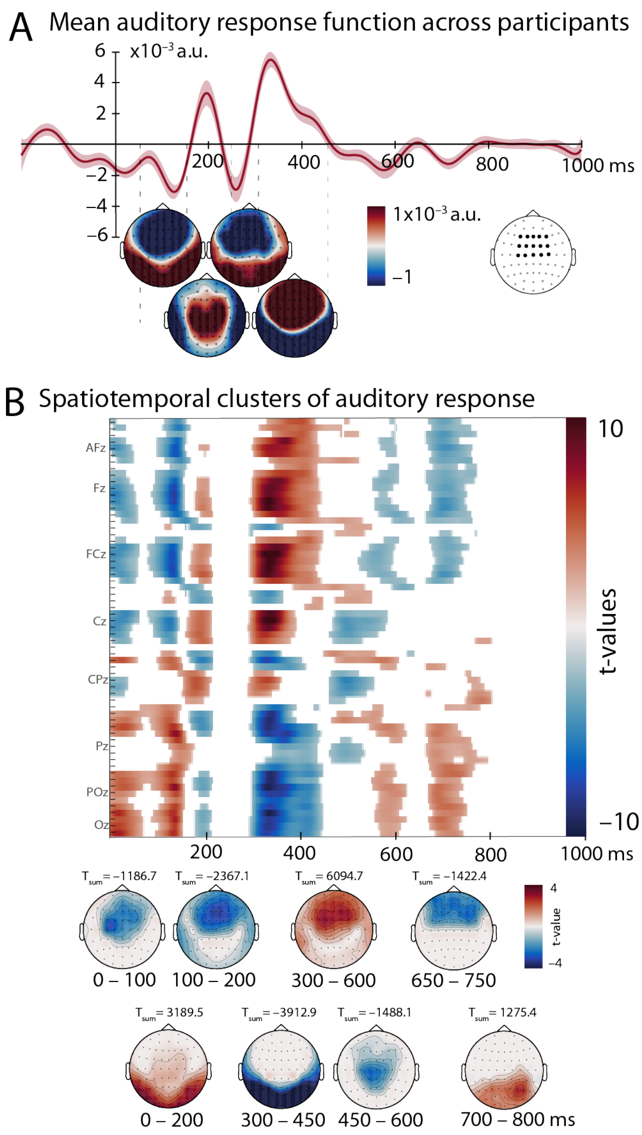


*Figure S2. Auditory response function (using motion and audio regressor simultaneously) for adult participants.* A) shows the generic model (mean ± within-subject SEM) computed across all participants, averaged over F3, F1, Fz, F2, F4, FC3, FC1, FCz, FC2, FC4, C3, C1, Cz, C2, and C4, and topographic representations for 50 – 150 ms, 150 – 250 ms, 250 – 300 ms, and 300 – 450 ms with electrodes included in the above-shown average marked by black dots. B) displays the results of the cluster-based permutation test, comparing the response function shown in A). Positive deviations are displayed in red, while negative deviations are shown in blue. In the bottom part of B), the same clusters as in the top part of B) are shown as topographic distributions, along with the summed t-value across the cluster.

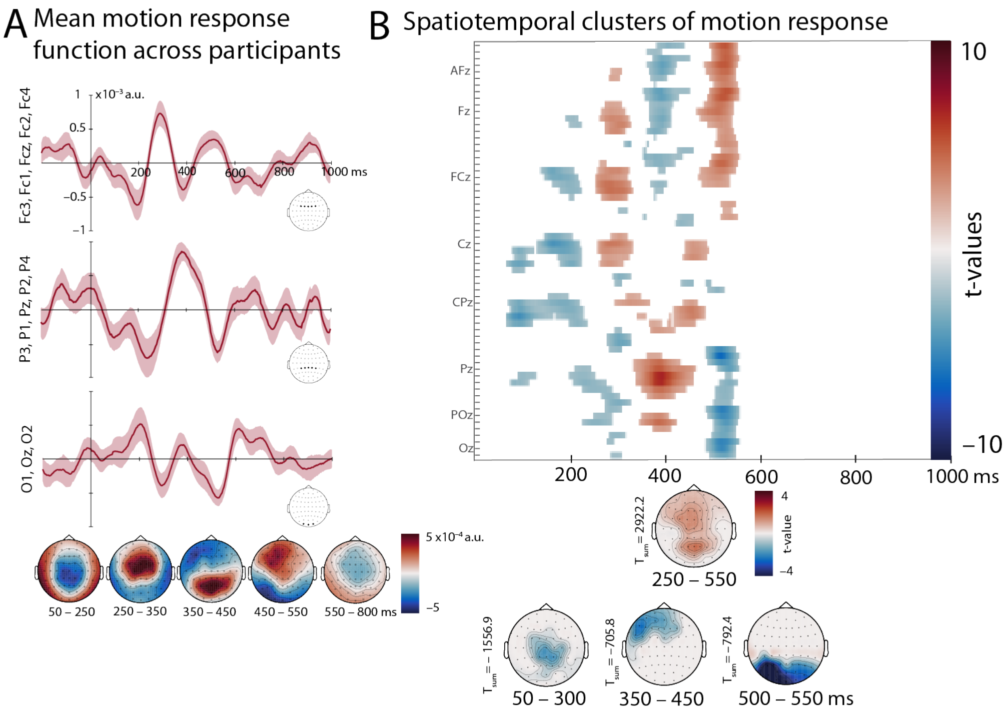


*Figure S3. Motion response function (using motion and audio regressor simultaneously) for adult participants.* A) shows the generic model (mean ± within-subject SEM) computed across all participants and averaged for three electrodes groups, frontocentral (top row), posterior (middle row), and occipital (bottom row). Topographic representations are shown for 50 – 250 ms, 250 – 350 ms, 350 – 450 ms, 450 – 550 ms, and 550 – 800 ms. B) displays the results of the cluster-based permutation test, comparing the response function shown in A) to zero. Positive deviations are displayed in red, while negative deviations are shown in blue. In the bottom part of B), the same clusters as in the top part of B) are shown as topographic distributions, along with the summed t-value across the cluster.

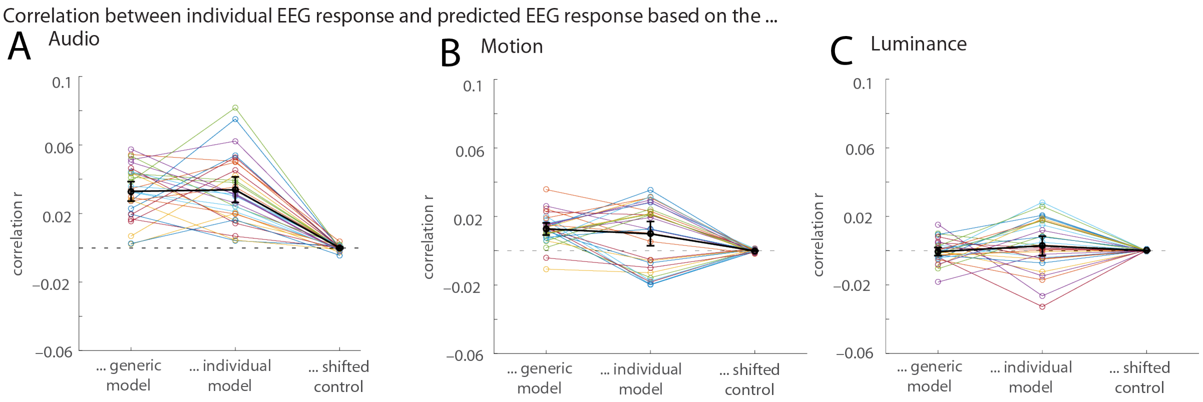


*Figure S4. Correlation between model and EEG response for adult participants.* The recorded individual EEG response was correlated with three different parameters using Pearson's correlation coefficient for the audio regressor (A), motion regressor (B), and luminance regressor (C). On the left, the correlation between the recorded EEG responses of participant n and the response predicted by the generic model based on the remaining n-1 participants is shown for each participant. In the middle, the correlation between the model trained on the first 80 % of the data available for each participant and used to predict the remaining 20 % from that participant and the actual EEG response recoded from that participant is shown. The right column shows the correlation between the prediction generated by the generic model and the recorded EEG data shifted in a circular way in steps of 2 s as a control condition (averaged over all possible shifts). Correlations are shown for each infant participant (in colors) as well as the mean correlation with 95% CI (confidence interval) across all participants (in black).

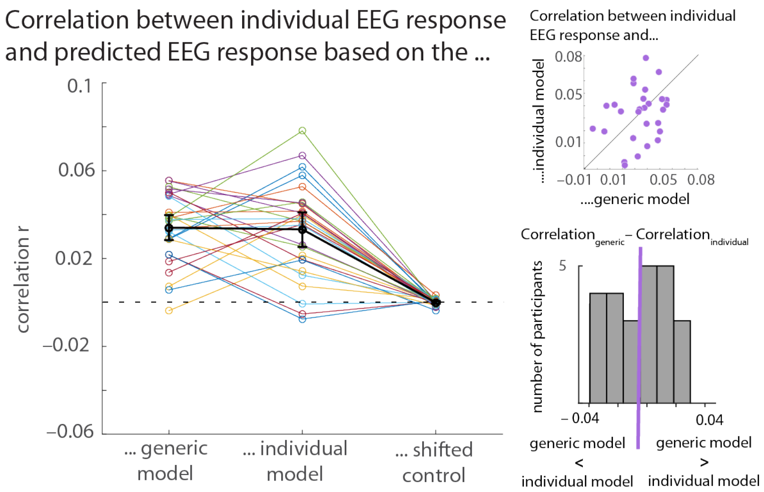


*Figure S5. Correlation between model and EEG response for adult participants using motion and audio regressors in one model.* The left part of the figure shows the correlation (based on Pearson's correlation coefficient) between the recorded EEG signal and the EEG responses predicted based on the generic model (left column), the individual model (middle column), and a shifted control condition (right column, see text). The two plots on the right hand visualize a comparison between the generic and the individual model. In the top plot, each purple dot indicates the difference between the correlation with the general model and the correlation with the individual model. Hence, a purple dot in the right bottom part of the graph indicates an individual with a higher correlation for the generic compared to the individual model, while a purple dot in the top left part indicates an individual with a higher correlation for the individual compared to the generic model. The bottom plot displays the same information in a bar graph; number of individual having a higher correlation for the generic model have a positive difference and hence fall to the right of the zero-threshold marked in purple while those with a higher correlation for the individual model have a negative difference and fall to the left of the zero-threshold.
